# Non-adhesive alginate hydrogels support growth of pluripotent stem cell-derived intestinal organoids

**DOI:** 10.1101/364885

**Authors:** Meghan M. Capeling, Michael Czerwinski, Sha Huang, Yu-Hwai Tsai, Angeline Wu, Melinda S. Nagy, Benjamin Juliar, Yang Song, Nambirajan Sundaram, Shuichi Takayama, Eben Alsberg, Michael Helmrath, Andrew J. Putnam, Jason R. Spence

**Author notes:** Author for correspondence JRS.

## Abstract

Human intestinal organoids (HIOs) represent a powerful system to study human development and are promising candidates for clinical translation as drug-screening tools or engineered tissue. Experimental control and clinical use of HIOs is limited by growth in expensive and poorly defined tumor-cell-derived extracellular matrices, prompting investigation of synthetic ECM-mimetics for HIO culture. Since HIOs possess an inner epithelium and outer mesenchyme, we hypothesized that adhesive cues provided by the matrix may be dispensable for HIO culture. Here, we demonstrate that alginate, a minimally supportive hydrogel with no inherent cell adhesion properties, supports HIO growth *in vitro* and leads to HIO epithelial differentiation that is virtually indistinguishable from Matrigel-grown HIOs. Additionally, alginate-grown HIOs mature to a similar degree as Matrigel-grown HIOs when transplanted *in vivo*, both resembling human fetal intestine. This work demonstrates that purely mechanical support from a simple-to-use and inexpensive hydrogel is sufficient to promote HIO survival and development.

## Introduction

A pivotal development in the fields of developmental biology and regenerative medicine was the use of human pluripotent stem cells (hPSCs) to generate human cell types, tissues, and organoid model systems *in vitro* (Huch et al., 2017; Johnston, 2015; Little, 2017). hPSC-derived organoids are 3-dimensional (3D) organ-like tissues that partially recapitulate structural and functional aspects of the organs after which they are modeled (Dye et al., 2016; Dye et al., 2015; Eiraku et al., 2011; Lancaster et al., 2013; Miller et al., 2018; Spence et al., 2011; Takasato et al., 2014; Takebe et al., 2013). In particular, human intestinal organoids (HIOs) are a useful tool to study intestinal development (Aurora and Spence, 2016; Dedhia et al., 2016; Finkbeiner et al., 2015a; Finkbeiner et al., 2015b; Munera et al., 2017; Singagoga and Wells, 2015; Tsai et al., 2016; Tsai et al., 2017), evaluate gut-microbe interactions (Hill et al., 2017), and model chronic health conditions such as Crohn’s disease and inflammatory bowel disease (Wells and Spence, 2014).

HIOs are generated from hPSCs through a step-wise differentiation process resulting in the formation of multi-cellular spheroids which are embedded in a 3D extracellular matrix (ECM) to support growth and development. Spheroids grow into HIOs over the course of approximately 30 days in culture (Spence et al., 2011). Currently, basement membrane-like ECM derived from mouse sarcoma cells, sold under brand names such as Matrigel or Cultrex, are standard for organoid culture. There are many issues with these cell-derived ECMs, including batch-to-batch variability, inability to control biophysical and biochemical properties, and a potential for pathogen transfer (Czerwinski and Spence, 2017). Perhaps most importantly, cell-derived ECMs typically have an uncharacterized protein composition and introduce a largely uncontrollable biological variability into experimental design. Lastly, cell-derived ECMs have a high cost, hindering scale-up. Collectively, these factors limit biological control during experiments and hamper downstream clinical applications. These limitations have prompted investigation into fully defined synthetic matrices to support organoid culture *in vitro*. Thus far only polyethylene glycol (PEG) has been utilized for HIO culture, motivating research into other hydrogel systems. Recent work has shown that modified PEG hydrogels can be engineered to mimic Matrigel to support HIO growth (Cruz-Acuna et al., 2017), or to support epithelium-only organoids (enteroids) generated from isolated murine or human intestinal stem cells (Gjorevski et al., 2016). These studies focused on engineering hydrogels to mimic the stiffness, adhesivity, and degradation dynamics required to support cellular viability and attachment (Cruz-Acuna et al., 2017; Gjorevski et al., 2016). However, given that hPSC-derived HIOs are comprised of both epithelium and an outer mesenchyme with supportive basement membrane, we hypothesized that HIOs may create their own niche and thus may be amenable to growth in substrates lacking inherent cell recognition.

In this work, we present evidence that native or unmodified alginate can be used as a simple hydrogel system that supports HIO growth and development *in vitro* and transplantation *in vivo* into immunocompromised mice. Alginate is an FDA approved polysaccharide derived from algae that is favorable due to its biocompatibility and ease of manipulation. Use of alginate does not require specialized bioengineering skills, and gelation is controlled by crosslinking with calcium; thus, alginate can be implemented using commercially available reagents without a need for further modification. Since unmodified alginate does not possess cell adhesive properties and its hydrophilic nature inhibits protein adsorption, it is a minimally supportive matrix that provides mechanical support for HIOs in a 3D environment. Here, we report that alginate hydrogels supported HIO viability and development *in vitro*. HIOs cultured in alginate grew optimally at a stiffness that was comparable to Matrigel, and the resulting alginate-grown HIOs were highly similar to Matrigel-grown HIOs *in vitro*. Alginate and Matrigel-grown HIOs underwent similar engraftment and maturation when transplanted *in vivo*, and both closely resembled human fetal intestine after transplantation. Collectively, these results demonstrate the effectiveness of alginate as a support matrix for HIOs and as an alternative for cell-derived ECM. Alginate overcomes many limitations of Matrigel and is significantly more cost effective than either Matrigel or PEG, making it a promising solution to improve experimental control and increase the clinical potential of organoids.

## Results

### Alginate hydrogels support HIO viability

We identified alginate as a potential HIO growth matrix based on its cost-effectiveness, biocompatibility, mild gelation conditions (Lee and Mooney, 2012), ability to control physical and biochemical properties (Jeon et al., 2013; Samorezov et al., 2015), and viscoelastic behavior (Webber and Shull, 2004). Many extracellular matrices and soft tissues exhibit viscoelastic behavior *in vivo* (Chaudhuri, 2017), including embryonic tissue (Forgacs et al., 1998). Previous work demonstrated that modified PEG hydrogels engineered to mimic the adhesive and biomechanical properties of Matrigel could support HIO development (Cruz-Acuna et al., 2017; Gjorevski et al., 2016); however, we hypothesized that due to the combined mesenchymal and epithelial composition of HIOs that establishes a laminin-rich basement membrane (Arora et al., 2017; Spence et al., 2011), a simple non-adhesive hydrogel with a similar stiffness to Matrigel may also support HIO growth. In order to design a simple, cost-effective system, we explored the potential of unmodified alginate to support HIO expansion.

After inducing hPSCs toward an intestinal lineage as previously described (Finkbeiner et al., 2015a; McCracken et al., 2011; Spence et al., 2011; Tsai et al., 2016; Tsai et al., 2017), 3D hindgut spheroids self-assemble and detach from the 2D monolayer. These spheroids develop into HIOs over the course of approximately 30 days in 3D culture. Spheroids were collected and embedded in Matrigel or in alginate hydrogels spanning a range of polymer densities (0.5%-4% wt/vol). Alginate solutions containing spheroids were ionically crosslinked with a calcium chloride solution to form 3D hydrogel networks. Since matrix mechanical properties have been shown to impact HIO viability (Cruz-Acuna et al., 2017) as well as epithelial cell behavior (Enemchukwu et al., 2016), we varied the properties of alginate hydrogels by varying polymer density in order to identify an optimal matrix to support cell viability (Fig. 1). Using rheometry, we measured the mechanical properties of alginate gels with an in situ gelation test. Rheological data confirm that both the storage and loss moduli of alginate hydrogels increase with increasing polymer density. The mechanical properties of Matrigel fall within the range of alginate hydrogels tested. Both the storage and loss moduli of 1% alginate were most similar to previously reported storage/loss moduli of Matrigel (Cruz-Acuna et al., 2017) (Fig. 1a).

**Figure 1.**
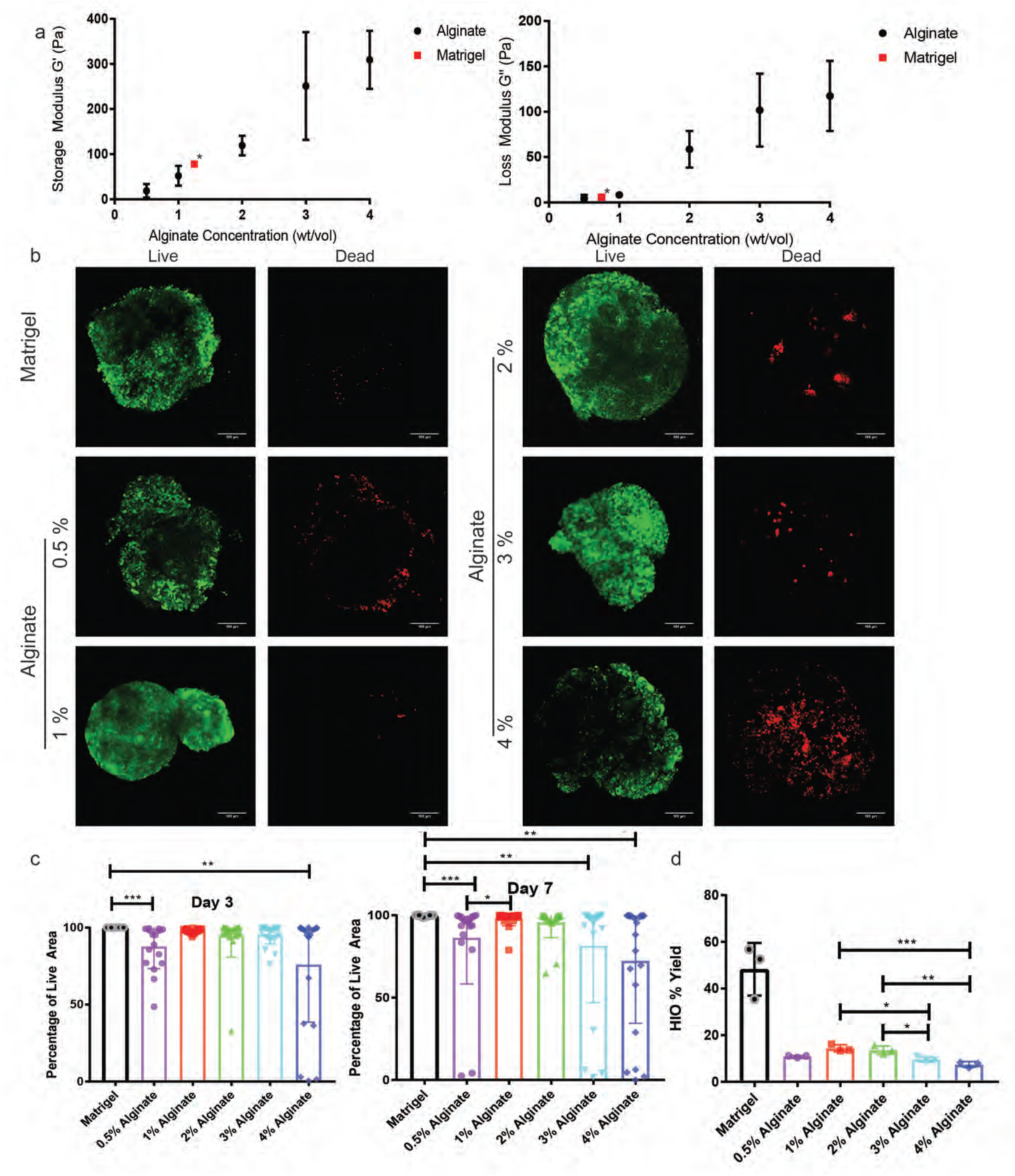
Alginate supports HIO survival *in vitro*. **(a)**: Rheological characterization of alginate hydrogels. Data shown are the mean ± standard deviation from n≥3 gels per condition. *Matrigel properties from previously published data (Cruz-Acuna et al., 2017). **(b)**: Representative images of live (Calcein AM, green) and dead (Ethidium homodimer-1, red) staining of spheroids in alginate and Matrigel after 7 days in culture. Scale bar = 100μm. **(c)**: Quantification of spheroid viability after 3 and 7 days encapsulated in alginate or Matrigel. Percentage of live area denotes area of spheroid expressing live marker over total spheroid area. Data are combined from 3 independent experiments with n>6 spheroids per condition per experiment. Each point depicts viability of an individual spheroid, while bars depict mean and standard error. Significance was calculated with a one-way ANOVA and multiple comparisons test. **(d)**: Quantification of HIO yield after 28 days in culture. HIO yield was calculated as the percentage of spheroids which gave rise to HIOs. Data shown are the average yields from 3 independent experiments with n>100 spheroids per condition. Each point depicts mean yield from one experiment, while bars depict mean and standard error. Significance was calculated with a one-way ANOVA and multiple comparisons test.

To optimize alginate growth conditions throughout the course of HIO development, we established metrics of success at both early and late time points. We examined spheroid viability at days 3 and 7 after encapsulation to assess the initial response of spheroids to alginate (Fig. 1b. c), as well as quantified overall HIO yield after 28 days, calculated as the percentage of embedded spheroids which matured into HIOs (Fig. 1d). Using live-dead staining, HIO viability was quantified as the percentage of spheroid area live-stained at days 3 and 7 post-encapsulation. Results presented in Figure 1c represent combined data from 3 independent experiments where n≥6 spheroids were analyzed at each condition. At 3 days post-encapsulation, none of the Matrigel spheroids (0%) displayed signs of death while spheroids in all concentrations of alginate displayed some cell death. After 3 days, spheroids in both 0.5% alginate and 4% alginate displayed significant decreases in viability, while spheroids in 1%, 2%, and 3% alginate displayed similar viability compared to spheroids in Matrigel (Fig. 1c). We noted a highly variable degree of viability at 4% alginate with some spheroids exhibiting high viability with others exhibiting complete death. By 7 days post-encapsulation, spheroids embedded in both 1% and 2% alginate retained the highest viability and HIOs grown in 0.5%, 3%, and 4% alginate exhibited decreased average viability from day 3 to day 7 with multiple spheroids showing 0% viability in each of these conditions (Fig. 1c). On the other hand, spheroids grown in 1% and 2% alginate remained nearly 100% viable with only a few spheroids displaying cell death. Thus, by day 7 it is apparent that spheroid survival depends on alginate density, with 1% and 2% alginate best supporting viability.

HIO yield was calculated as the percentage of spheroids at day 1 of encapsulation that gave rise to HIOs after 28 days (Fig. 1d). All of the alginate concentrations tested produced significantly lower HIO yields than Matrigel, although we note that our yield in Matrigel is 3.7 fold higher than those reported by others (Arora et al., 2017). The low HIO yields at day 28 as compared with high viability in alginate at day 7 suggests that additional death may occur later in HIO development, or that certain spheroids do not expand in alginate. HIO yield was highest in 1% alginate as compared with all other alginate formulations tested, although there was no significant difference in yield between 0.5%, 1%, and 2% alginate. We selected 1% or 2% alginate as the optimal concentrations for future experiments since these concentrations resulted in the highest spheroid viability at early time points as well as highest overall HIO yield.

### The epithelium of alginate-grown HIOs and Matrigel-grown HIOs is indistinguishable

After 28 days of culture, we used histological techniques to compare HIOs cultured in alginate and Matrigel. Histological analysis with hematoxylin and eosin (H&E) staining revealed that HIOs cultured in both alginate and Matrigel form an inner epithelium surrounded by an outer mesenchyme (Fig. 2a). While the epithelium of alginate and Matrigel-grown HIOs appears quite similar histologically, the mesenchyme of Matrigel-grown HIOs invades the surrounding matrix whereas alginate-grown HIOs do not appear to invade the hydrogel.

**Figure 2.**
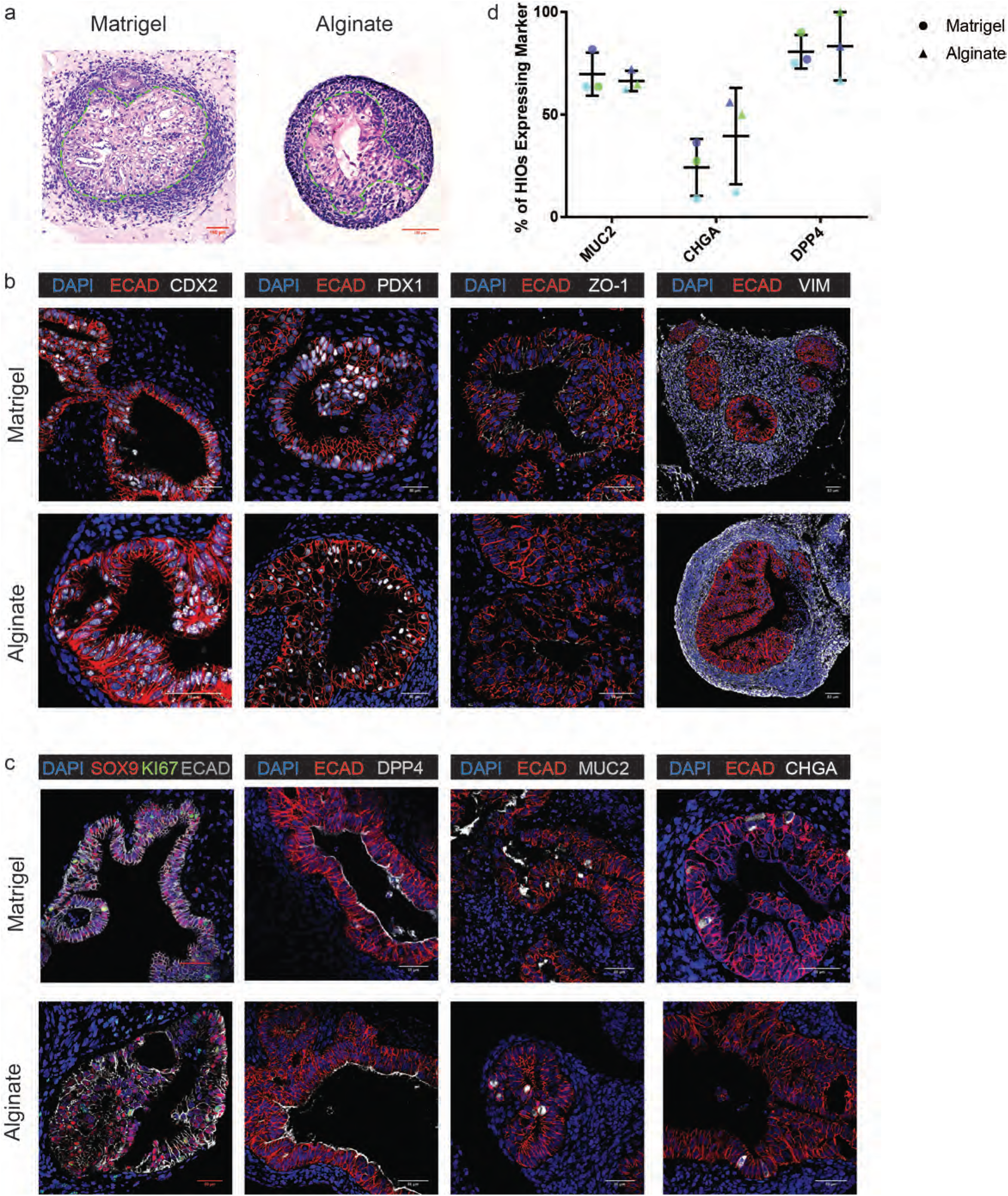
Epithelium of alginate HIOs resembles epithelium of Matrigel HIOs *in vitro*. **(a)**: Hematoxylin and eosin staining of alginate and Matrigel HIOs cultured in 1% alginate and Matrigel for 28 days. Dashed lines outline the epithelium. **(b)**: Representative images of general epithelial marker staining in HIOs cultured in 1% alginate and Matrigel for 28 days. Markers shown are ECAD (epithelial marker), CDX2 (intestinal epithelium marker), PDX1 (duodenum marker), ZO-1(tight junction marker), and VIM (mesenchymal marker). Scale bar = 50μm. **(c)**: Representative images of specific epithelial cell marker staining in HIOs cultured in 1% alginate and Matrigel for 28 days. Markers shown are SOX9 (progenitor cell marker), KI67 (proliferative cell marker), DPP4 (small intestinal brush border enzyme marker), MUC2 (goblet cell marker), and CHGA (enteroendocrine marker). Scale bar = 50μm. **(d)**: Frequency of mature cell type differentiation in 1% alginate and Matrigel. Points depict the percentage of HIOs expressing each marker for 3 independent experiments with n≥6 HIOs per experiment. Each color represents one matched experiment. Significance was calculated with a one-way ANOVA and multiple comparisons test.

HIOs cultured in both alginate and Matrigel were examined for the presence of markers of intestinal epithelial patterning, mesenchyme formation, and polarization (Fig. 2b) as well as markers of fully differentiated intestinal cell types including enterocytes, goblet cells, and enteroendocrine cells (Fig. 2c). The epithelium of both alginate and Matrigel-grown HIOs expressed the intestinal epithelial marker CDX2, confirming that alginate-grown HIOs differentiate along an intestinal lineage as observed in Matrigel-grown HIOs. Additionally, both alginate and Matrigel-grown HIOs expressed the duodenum marker PDX1 throughout the epithelium, indicating that HIOs cultured in both matrices became patterned into proximal small intestine (Tsai et al., 2017). Alginate and Matrigel-grown HIOs expressed the tight junction marker ZO-1 at apical surfaces suggesting proper epithelial polarization. Additionally, both alginate and Matrigel-grown HIOs supported the development of an outer mesenchyme as evidenced by the presence of VIM expression surrounding the epithelium.

SOX9 is expressed in progenitor cells in HIOs (Hill et al., 2017), and we observed that the majority of epithelial cells in both alginate and Matrigel-grown HIO expressed SOX9 after 28 days of culture *in vitro*. Co-staining with the proliferation marker KI67 indicates that both alginate and Matrigel-grown HIOs are proliferative (Fig. 2c). Additionally, both alginate and Matrigel-grown HIOs gave rise to differentiated epithelial cell types *in vitro* as HIOs cultured in the two matrices displayed epithelial cells expressing markers for small intestinal enterocytes (DPP4), goblet cells (MUC2), and enteroendocrine cells (CHGA). This suggests that differentiation within alginate-grown HIOs develops along a similar timeline when compared to Matrigel-grown HIOs.

Although we observed differentiated cell markers within HIOs, the abundance of differentiation was generally low. To report the frequency of differentiation across multiple HIOs and across different batches of HIOs, we quantified the percentage of HIOs expressing the markers DPP4, MUC2, and CHGA across 3 independent experiments (Fig. 2d). Our results demonstrated that alginate and Matrigel-grown HIOs expressed differentiation markers with a similar frequency; however, we noted variability across batches of HIOs. Our data suggests that batch-to-batch variability is the most likely explanation for expression differences between experiments, and this batch effect is seen in alginate and Matrigel-grown HIOs. Taken together, these results demonstrate that alginate-grown HIOs resemble Matrigel-grown HIOs at the epithelial level and suggest that alginate is an effective alternative to Matrigel for supporting HIO development *in vitro*.

### Transplanted alginate HIOs mature *in vivo*

We next explored the engraftment and maturation potential of alginate-grown HIOs *in vivo*. Matrigel-grown HIOs and HIOs cultured in synthetic matrices have been shown to develop into more mature intestinal tissue after transplantation into immunocompromised mice (Cruz-Acuna et al., 2017; Finkbeiner et al., 2015a; Finkbeiner et al., 2015b; Watson et al., 2014), so a key success criterion of alginate HIOs is the ability to undergo similar maturation following transplantation into immunocompromised (NSG) mice. HIOs grown in both 2% alginate (n=1 mice) and Matrigel (n=12 mice) were dissociated from their growth matrices and implanted beneath the kidney capsules of NSG mice (Fig. 3a). After 10 weeks *in vivo*, transplanted HIOs (tHIOs) previously cultured in alginate developed crypt-villus structures with associated submucosa, lamina propria, and muscularis mucosae resulting in a comparable architecture to tHIOs previously cultured in Matrigel (H&E staining). Both alginate and Matrigel tHIOs displayed goblet cells throughout the villi (Alcian blue/PAS staining) as well as organized collagen fibers in the submucosa (Trichrome staining) (Fig. 3b). The structure and features of alginate and Matrigel tHIOs both closely resemble human fetal intestinal tissue (Fig. 3b). The epithelium of alginate tHIOs demonstrated evidence that they retained intestinal lineage identity (CDX2+) and remained patterned into duodenum (PDX1+) following transplantation (Fig. 3c). Alginate and Matrigel tHIOs both formed ZO-1+ tight junctions at the apical surface of the epithelium, similar to fetal intestinal tissue. Additionally, both alginate and Matrigel tHIOs demonstrated the expected localization of mesenchyme inside and below villi as demonstrated by VIM staining.

**Figure 3.**
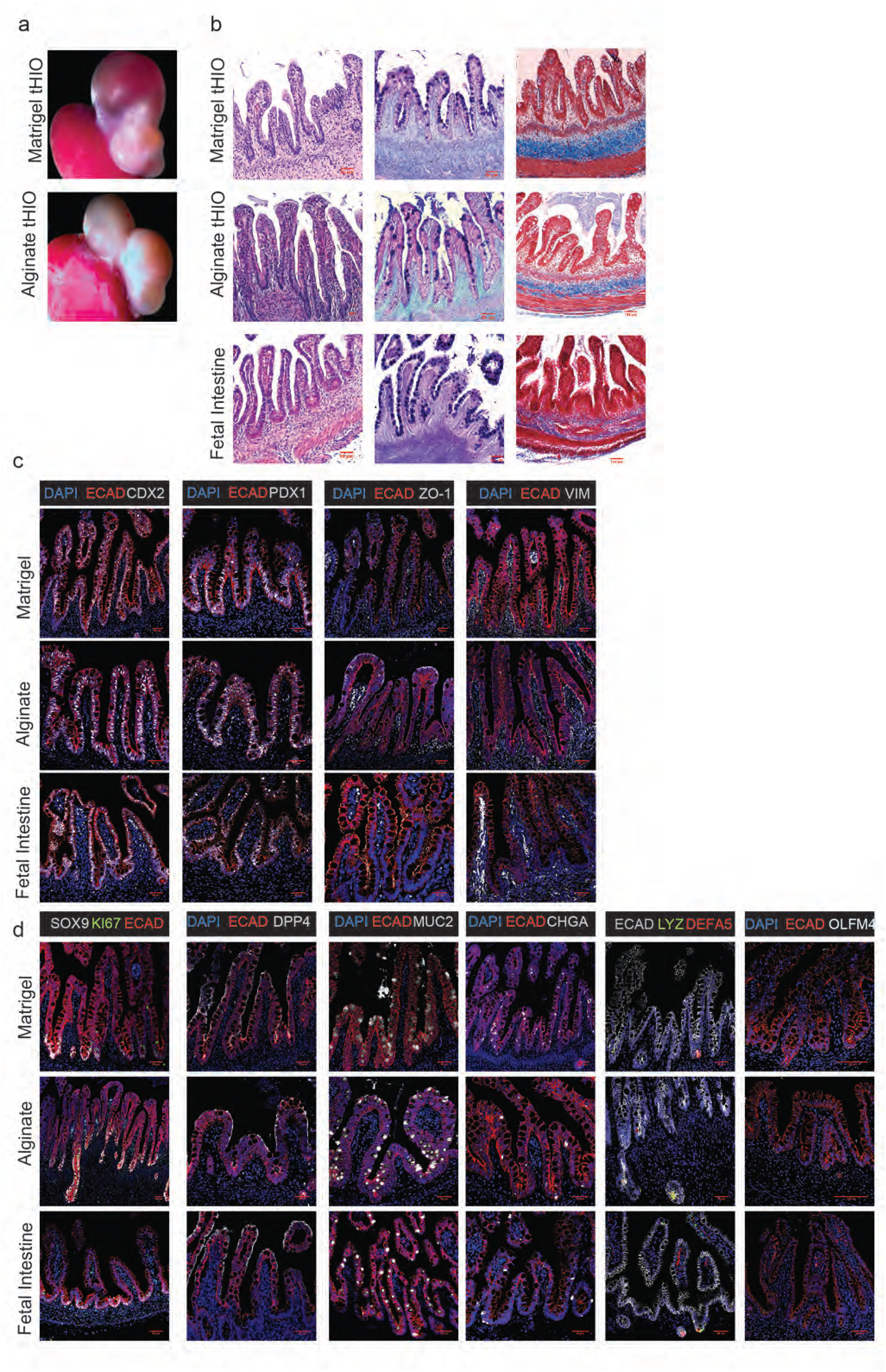
Alginate HIOs mature in similar manner as Matrigel HIOs *in vivo*. **(a)**: Dissected kidneys containing tHIOs cultured in alginate or Matrigel. HIOs were cultured in either Matrigel or 2% alginate dissolved in H_2_O and transplanted after 28 days of culture *in vitro*. **(b)**: Hematoxylin and eosin, Alcian blue/PAS, and Trichrome staining reveal the presence of mature crypt-villus structures in alginate and Matrigel tHIOs. Scale bar = 50μm (H&E and Alcian blue/PAS, 100 μm Trichrome. **(c)**: Representative images of general epithelial marker staining in tHIOs from and Matrigel. Markers shown are ECAD (epithelial marker), CDX2 (intestinal epithelium marker), PDX1 (duodenum marker), and VIM (mesenchymal marker). Scale bar = 50μm. **(d)**: Representative images of specific epithelial cell marker staining in tHIOs from alginate and Matrigel. Markers shown are SOX9 (progenitor cell marker), KI67 (proliferative cell marker), DPP4 (small intestinal brush border enzyme marker), MUC2 (goblet cell marker), CHGA (enteroendocrine cell marker), LYZ and DEFA5 (Paneth cell markers), and OLFM4 (intestinal stem cell marker). Two transplant experiments were conducted with a total of n= 11 Matrigel transplanted and n=12 alginate transplanted mice. Scale bar = 50μm for all markers except OLFM4 scale bar = 100μm.

After growth *in vivo*, alginate tHIOs increased in maturity as evidenced by lack of expression of SOX9 throughout the villi and increased expression of markers for differentiated cell types (Fig 3d). Both SOX9 and KI67 became localized to the crypt-like domains of alginate and Matrigel tHIOs as expected, suggesting the development of a mature crypt-villus axis with proliferative progenitor cells residing in the crypts. The proper localization of intestinal stem cells to the crypts in both alginate and Matrigel tHIOs was confirmed by OLFM4 expression (Dame et al., 2018). Alginate and Matrigel tHIOs similarly expressed DPP4, MUC2, and CHGA throughout the epithelium. Additionally, both alginate and Matrigel tHIOs supported the differentiation of Paneth cells localized to crypts as evidenced by co-expression of LYZ and DEFA5, consistent with previous reports (Finkbeiner et al., 2015b; Watson et al., 2014). Staining patterns in alginate and Matrigel tHIOs for all markers closely resembled staining patterns in human fetal intestine. Taken together these results demonstrate that HIOs grown in alginate differentiate and mature *in vivo* to a similar degree as Matrigel HIOs and highlights alginate as a viable alternative to Matrigel for HIO culture.

### The epithelium of alginate and Matrigel-grown HIOs share a high degree of molecular similarity *in vitro* and *in vivo*

We utilized RNA sequencing analysis to determine the degree of similarity or difference between alginate and Matrigel-grown HIO epithelia in an unbiased manner. In order to reduce variance across samples and specifically assess epithelial gene expression, we isolated the epithelium from alginate and Matrigel-grown HIOs and tHIOs, further expanded this epithelium in identical culture conditions, and then performed bulk RNA-sequencing (Fig. 4a). The epithelium from *in vitro* grown HIOs was isolated to create alginate and Matrigel HIO-derived epithelium-only (HdE) cultures. After 10 weeks *in vivo*, the epithelium from alginate and Matrigel tHIOs was similarly isolated and cultured *in vitro* to create alginate and Matrigel transplanted HIO-derived epithelium (tHdE) (Fig. 4a). Following RNA-sequencing, the transcriptomic data for individual replicates from all 4 groups were analyzed for their similarity using Pearson’s correlation coefficient. Alginate and Matrigel HdEs clustered together with high correlation, and similarly alginate and Matrigel tHdEs formed a cluster with high correlation. These clusters suggest that differences between culture conditions (*in vitro* vs. transplantation) are the main drivers of variability between samples as enteroids from *in vitro* grown alginate and Matrigel HIOs are more similar to each other than they are to their transplant-derived counterparts even when grown in uniform culture conditions. This is consistent with previous data showing that HIOs grown *in vitro* are immature/fetal in nature, and transplanted HIOs become more mature/adult-like (Hill et al., 2017).

**Figure 4.**
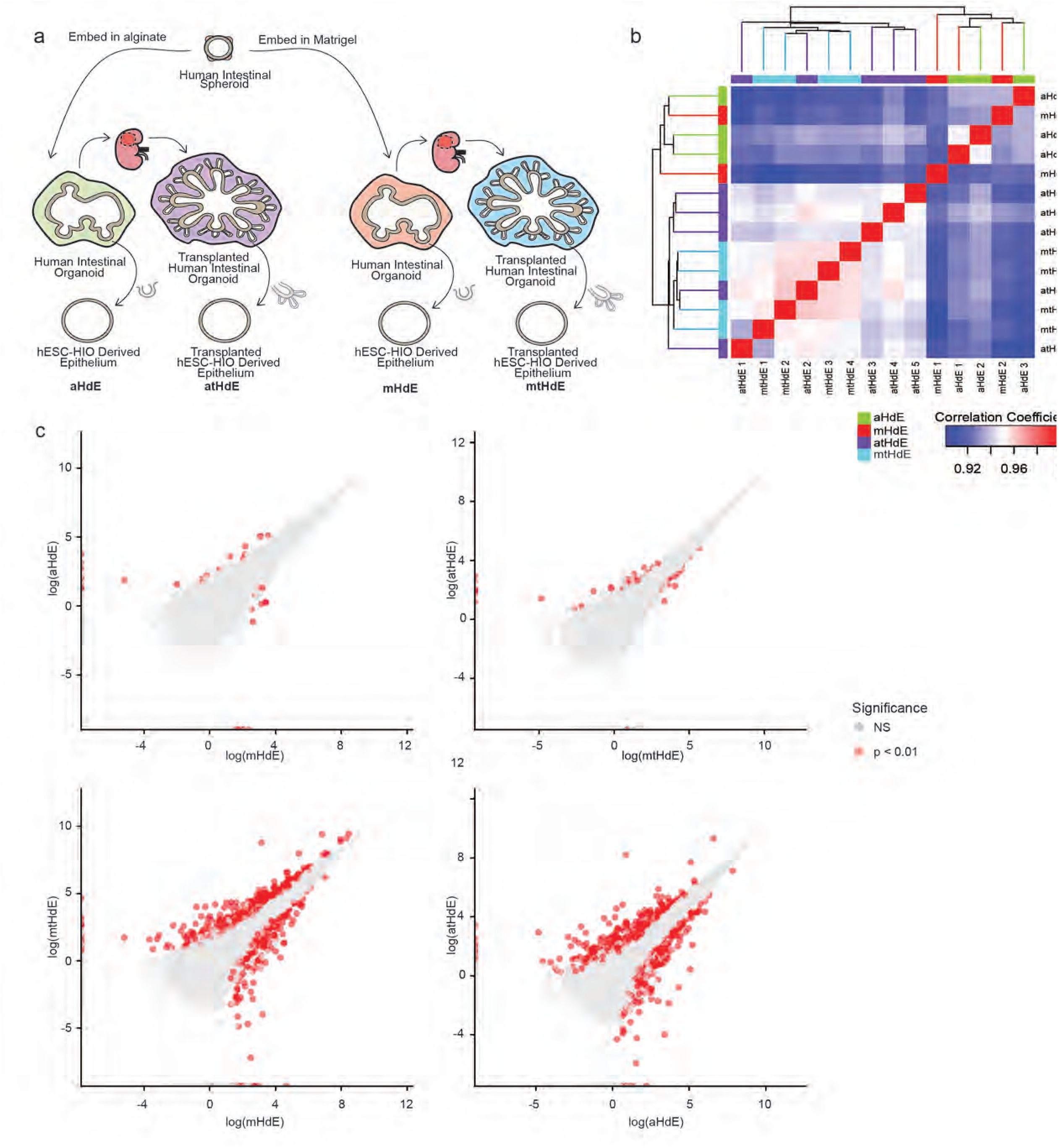
RNA-seq comparison of alginate and Matrigel HIO epithelia. **(a)**: Schematic overview of sample groups included in RNA-seq analysis. Epithelia were extracted from alginate and Matrigel HIOs both after culture *in vitro* and after transplantation into mice. **(b)**: Clustering HdEs and tHdEs by sample similarity using Pearson’s correlation coefficient (n=3 aHdE, n=2 mHdE, n=5atHdE, n=4mtHdE). Clusters formed between aHdEs and mHdEs as well as between atHdEs and mtHdEs. **(c)**: Differential expression analysis comparing aHdEs to mHdEs, atHdEs to mtHdEs, mHdEs to mtHdEs, and aHdEs to atHdEs. Red dots represent genes with significant differences in expression (p < 0.01). In these plots, the alginate and Matrigel samples are nearly identical.

Additionally, we utilized differential expression analysis to compare alginate and Matrigel HdEs and tHdEs (Fig. 4c). There was a high degree of similarity between alginate HdEs (aHdEs) and Matrigel HdEs (mHdEs) with only 32 genes (2.3%) showing significant differences in expression. Similarly, there was high similarity between alginate transplant-derived HdEs (atHdEs) and Matrigel transplant-derived HdEs (mtHdEs) with only 42 genes (3%) displaying significant differences in expression. In contrast, comparison of aHdEs to atHdEs and mHdEs to mtHdEs revealed 730 (51.7%) and 908 (64.4%) genes, respectively, with significant differences in expression. This analysis confirms that the epithelium of alginate HIOs is nearly identical to Matrigel HIOs both *in vitro* and *in vivo*.

## Discussion

In this work we identified alginate as a minimally supportive growth matrix which supports HIO development both *in vitro* and *in vivo*. We showed that HIO survival is dependent upon alginate polymer density, with concentrations of 1%-2% alginate (wt/vol) selected to best support viability at early time points as well as maximize overall HIO yield. While alginate supported HIO survival over at least 28 days in culture, the resulting yields were significantly lower than yields in Matrigel. However, the yields we observed in 1% and 2% alginate are comparable with yields previously reported for HIOs cultured in Matrigel, and could likely be optimized by sorting spheroids for size or characteristic to improve yield (Arora et al., 2017). It is unclear whether alginate results in lower overall yield of HIOs as compared with previously described synthetic polymers since viability was only reported at early time points in previous work (Cruz-Acuna et al., 2017). Lower HIO yields in alginate as compared to Matrigel may be due to the inability of cells to remodel the hydrogel, lack of interactions with serum proteins, or the lack of growth factors present in Matrigel.

HIOs cultured in alginate resulted in tissue indistinguishable from Matrigel HIOs at the epithelial level. Importantly, both matrices supported further differentiation and maturation *in vivo*, illustrating that HIOs cultured in alginate retain the potential to develop into mature intestinal tissue that resembles human fetal intestine. While the epithelia from alginate and Matrigel-grown HIOs are highly similar, it remains unclear whether or not there are differences at the mesenchymal level. It is interesting to note that the mesenchyme of alginate-grown HIOs did not invade the surrounding matrix as in Matrigel-grown HIOs. Further research is necessary to elucidate potential mesenchymal differences between alginate and Matrigel-grown HIOs.

Given the similarities between alginate and Matrigel-grown HIOs, alginate is an effective alternative to Matrigel-based culture systems which eliminates reliance on animal-derived materials and reduces cost, thereby increasing translational potential. The alginate utilized in our experiments costs approximately 320 times less than PEG and 700-900 times less than Matrigel (~$0.44 alginate vs. ~$140 PEG vs. ~$300-$400 Matrigel per 10 mL, depending on type), presenting a critical cost advantage for both basic and translational studies.

From a biological standpoint, perhaps the most interesting observation of HIO culture in alginate is that HIOs do not require external cues from the extracellular matrix. The alginate we utilized in this work was not modified with adhesive peptides to support HIO growth and was thus biologically inert, providing purely mechanical support to developing HIOs. The lack of adhesive or biochemical cues from the hydrogel suggests that HIOs are able to create their own niche, likely through the basement membrane and trophic support that is established between the epithelium and mesenchyme. The lack of chemical modifications makes alginate a simple system which lends nicely toward large scale production, but the polymer can easily be modified with such groups for further research (Augst et al., 2006; Rowley et al., 1999). However, as a natural matrix, alginate does come with the limitation of batch-to-batch variability as with Matrigel (Fu et al., 2010), but our experiments found that alginate was able to optimally support growth over a range of conditions, so small variations between batches should not cause a large change in matrix efficacy. The system described here can likely be implemented to support additional 3D culture systems in a simple, cost-effective manner to advance regenerative medicine.

## Experimental Procedures

### hESC lines and generation hPSC-Derived Intestinal Organoids

Human ES line H9 (NIH registry #0062) was obtained from the WiCell Research Institute. All experiments using human ES cells were approved by the University of Michigan Human Pluripotent Stem Cell Research Oversight Committee. ES cell lines are routinely karyotyped to ensure normal karyotype and monthly mycoplasma monitoring is conducted on all cell lines using the MycoAlert Mycoplasma Detection Kit (Lonza). H9 cells were authenticated using Short Tandem Repeat (STR) DNA profiling (Matsuo et al., 1999) at the University of Michigan DNA Sequencing Core and exhibited an STR profile identical to the STR characteristics published by Josephson et al. (Josephson et al., 2006). Stem cell maintenance and generation of hindgut spheroids was performed as described previously (Hill et al., 2017; Spence et al., 2011; Tsai et al., 2017). Briefly, hESC lines were induced to differentiate into endoderm using Activin A (100ng/mL, R & D Systems) for 3 days in RPMI1640 media supplemented with 0%, 0.2%, 2% HyClone dFBS on subsequent days. Endoderm was induced to differentiate into the intestinal lineage by treating cells for 5-6 days with FGF4 (500ng/mL, generated as previously described (Leslie et al., 2015)) and CHIR99021 (2μM). Following differentiation, free-floating hindgut spheroids were collected from differentiated stem cell cultures after days 5 and 6 of hindgut specification and plated in Matrigel (diluted with Advanced DMEM/F12 to a final concentration of 8 mg/mL) or alginate droplets as described below on a 24-well tissue culture grade plate. Organoid growth media consisted of Advanced DMEM/F12 supplemented with 1× B27 (Thermo Fisher, Waltham, MA), epidermal growth factor (EGF) (R&D Systems; 100 ng/mL), Noggin-Fc (100ng/mL) (purified from conditioned media (Heijmans et al., 2013)), and R-Spondin2 (5% conditioned medium (Bell et al., 2008)). Media was changed every 5-7 days.

### Epithelial Isolation

Generation of HdEs and tHdEs from HIOs and tHIOs was performed using previously described methods to isolate the epithelium (Tsai et al., 2018). In summary, HIOs and tHIOs were incubated in dispase (07923; STEMCELL Technologies) for 30 minutes on ice. Following incubation, dispase was removed and replaced with 100% fetal bovine serum for 15 minutes on ice. To mechanically separate the epithelium from mesenchyme, a volume of advanced Dulbecco’s modified Eagle medium/F12 (12634010; Gibco) equal to the initial volume of fetal bovine serum was added to the tissue before vigorously pipetting the mixture several times. Epithelial fragments then settled to the bottom where they were collected manually on a stereoscope by pipet. The epithelium was washed with ice-cold advanced Dulbecco’s modified Eagle medium/F12 and allowed to settle to the bottom of a 1.5-mL Eppendorf tube. The media was then withdrawn from the loose tissue pellet and replaced with Matrigel on ice. The Matrigel containing the isolated epithelium was gently mixed to evenly suspend the cells before being pipetted into individual 50-μL droplets in a 24-well plate. The plate containing the droplets was incubated at 37°C for 15 minutes to allow the Matrigel to solidify before adding LWRN growth media containing Thiazovivin (2.5 μmol/L), SB431542 (100 nmol/L), CHIR99021 (4 μmol/L), and Y27632 (10 μmol/L). After 24 hours, the media was replaced with LWRN growth media containing TZV (2.5 μmol/L), SB431542 (100 nmol/L), and CHIR99021 (4 μmol/L). After 3 days, cultures were maintained with LWRN growth media replaced every other day.

### Human Tissue

Normal, de-identified human fetal intestinal tissue was obtained from the University of Washington Laboratory of Developmental Biology. Tissue sections were obtained from formalin fixed, paraffin embedded 14-15 week fetal intestinal specimens. All research utilizing human tissue was approved by the University of Michigan institutional review board.

### Mouse Kidney Capsule Transplantation

The University of Michigan and Cincinnati Children’s Hospital Institutional Animal Care and Use Committees approved all animal research. Prior to transplantation, HIOs were mechanically dissociated from either alginate or Matrigel. HIOs were implanted under the kidney capsules of immunocompromised NOD-scid IL2Rg-null (NSG) mice (Jackson Laboratory strain no. 0005557) as previously described (Finkbeiner et al., 2015b). In summary, mice were anaesthetized using 2% isoflurane. A left-flank incision was used to expose the kidney after shaving and sterilization with isopropyl alcohol. HIOs cultured in alginate and Matrigel were surgically implanted beneath mouse kidney capsules using forceps. Prior to closure, an intraperitoneal flush of Zosyn (100 mg kg^−1^; Pfizer) was administered. Mice were euthanized for retrieval of tHIOs after 10 weeks. Results shown are representative of two experiments performed with a total of n=11 mice (Matrigel tHIOs) and n=12 mice (alginate tHIOs), with at least one organoid implanted per kidney capsule, depending on HIO size.

### Alginate Gel Formation

Low viscosity sodium alginate powder (Alfa Aesar, B25266) was dissolved in 1 mL of 1 × PBS or H_2_O to dilutions of 0.5%-4% (wt/vol). The alginate solution was then heated to 98°C for 30 minutes on a heating block to improve sterility and ensure that the alginate fully dissolved. Spheroids were suspended in the alginate solutions at a density of approximately 50 spheroids per 45 μL. 5 μL droplets of 2% calcium chloride (wt/vol) were deposited on the bottom of 24-well tissue culture plates (Nunclon). 45 μL of alginate containing spheroids was then pipetted directly onto the calcium solution to initiate ionic crosslinking, which began instantaneously upon pipetting. The gels were placed into a tissue culture incubator and allowed to fully polymerize for 20 minutes at 37°C before media was added. HIOs cultured in 1% and 2% alginate were used to obtain all data in Figures 2-4.

### Mechanical Characterization of Alginate Hydrogels

The storage and loss moduli of the alginate gels were determined by performing *in situ* gelation tests on an AR-G2 rheometer equipped with a peltier stage and 20 mm measurement head. In brief, 90μL of alginate was deposited onto the bottom of the rheometer while 10μL of 2% CaCl_2_ was deposited onto the measurement head. The head was lowered to a gap height of 300μm initiating gelation upon contact, and then the edges of the gel were sealed with mineral oil. The mechanical response of the gels was recorded by performing time sweep measurements at a constant strain of 6% and a frequency of 1 rad/s. The time sweep was continued until storage and loss moduli reached steady state indicating completion of gelation.

### Viability Assay and Quantification

Alginate gels and Matrigel droplets were incubated in 1 μM Calcein-AM (live; ThermoFisher), and 0.5 μM Ethidium-homodimer (dead; ThermoFisher) in PBS for 30 minutes. Samples were imaged using an Olympus IX73 Inverted microscope or Nikon A-1 confocal microscope. Quantification of viability was performed by calculating the percentage of the total projected area of a spheroid/organoid that stained positive for the live or dead stain using ImageJ (National Institute of Health, USA). The results are representative of three independent experiments performed with n≥6 gel samples per experimental group.

### RNA-sequencing and Bioinformatics Analysis

RNA isolation and analysis was carried out as previously described (Tsai et al., 2018). In summary, RNA from each sample was isolated using MagMAX-96 Total RNA (AM1830; Applied Biosystems) RNA isolation kits and used as input for library generation with Takara SMARTer Stranded Total RNA Sample Prep Kit (634876; Takara Bio USA). Samples were sequenced for 50-bp single-end reads across 10 lanes on an Illumina HiSeq 2500 by the University of Michigan DNA Sequencing Core. All reads were quantified using Kallisto pseudo alignment to an index of transcripts from all human genes within the Ensembl GRCh38 database (Bray et al., 2016). Gene level data generated from Kallisto was used to create normalized data matrix of pseudoaligned sequences (Transcripts Per Million, TPM) and differential expression was calculated using the Bioconductor package DEseq2. Estimated counts per transcript using the Bioconductor package tximport. Differential expression analysis was performed using the Bioconductor package DESeq2 using gene count data (Love et al., 2014). A gene was considered to be differentially expressed if it had a 2-fold or larger difference between groups and an adjusted *P* value of .01 or less. Principal component analysis and sample clustering were performed in R with log2 transformed and centered gene counts of gene level data on all genes that had a sum of at least 10 counts across all samples. Replicates for all samples were clustered by Euclidian distance, and pairwise Pearson correlation coefficients were plotted in R. All reads are deposited at the EMBL-EBI ArrayExpress archive under accession E-MTAB-7000.

### Tissue preparation, Immunohistochemistry, Electron Microscopy and imaging

#### Paraffin sectioning and staining

HIO and tHIO tissues were fixed in 4% Paraformaldehyde (Sigma) overnight and then dehydrated in an alcohol series: 30 minutes each in 25%, 50%, 75% Methanol:PBS/0.05% Tween-20, followed by 100% Methanol, and then 100% Ethanol. Tissue was processed into paraffin using an automated tissue processor (Leica ASP300). Paraffin blocks were sectioned 7 uM thick, and immunohistochemical staining was performed as previously described (Spence et al., 2009). A list of antibody information and concentrations used can be found in Table 1. H&E staining was performed using Harris Modified Hematoxylin (FisherScientific) and Shandon Eosin Y (ThermoScientific) according to manufacturing. PAS Alcian blue staining was performed using the Newcomer supply Alcian Blue/PAS Stain kit (Newcomer Supply, Inc.) according to manufacturer’s instructions. Trichrome staining was performed by the University of Michigan *in vivo* Animal Core.

**Table 1.**
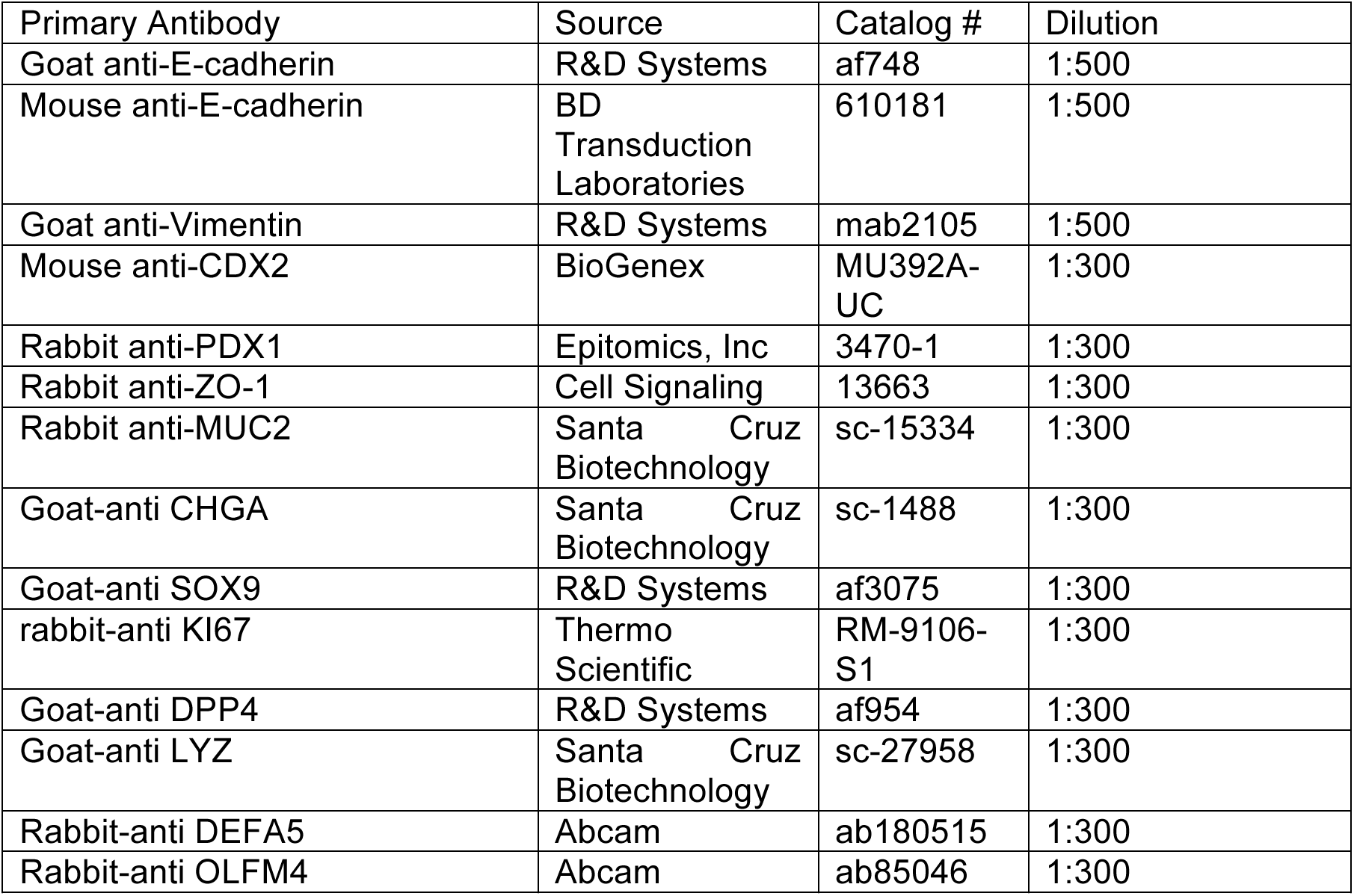
Antibody Information.

| Primary Antibody | Source | Catalog # | Dilution |
| --- | --- | --- | --- |
| Goat anti-E-cadherin | R&D Systems | af748 | 1:500 |
| Mouse anti-E-cadherin | BD<br>Transduction<br>Laboratories | 610181 | 1:500 |
| Goat anti-Vimentin | R&D Systems | mab2105 | 1:500 |
| Mouse anti-CDX2 | BioGenex | MU392A-UC | 1:300 |
| Rabbit anti-PDX1 | Epitomics, Inc | 3470-1 | 1:300 |
| Rabbit anti-ZO-1 | Cell Signaling | 13663 | 1:300 |
| Rabbit anti-MUC2 | Santa Cruz<br>Biotechnology | sc-15334 | 1:300 |
| Goat-anti CHGA | Santa Cruz<br>Biotechnology | sc-1488 | 1:300 |
| Goat-anti SOX9 | R&D Systems | af3075 | 1:300 |
| rabbit-anti Ki67 | Thermo<br>Scientific | RM-9106-S1 | 1:300 |
| Goat-anti DPP4 | R&D Systems | af954 | 1:300 |
| Goat-anti LYZ | Santa Cruz<br>Biotechnology | sc-27958 | 1:300 |
| Rabbit-anti DEFA5 | Abcam | ab180515 | 1:300 |
| Rabbit-anti OLFM4 | Abcam | ab85046 | 1:300 |

#### Imaging and image processing

Fluorescently stained slides were imaged on a Nikon A-1 confocal microscope. Brightness and contrast adjustments were carried out using ImageJ (National Institute of Health, USA) and adjustments were made uniformly across images.

### Quantification and Statistical Analysis

Statistical analyses and plots were generated in Prism 6 software (GraphPad). If more than two groups were being compared within a single experiment, an unpaired one-way ANOVA was performed followed by Tukey’s multiple comparisons test to compare the mean of each group with the mean of every other group within the experiment. For all statistical tests, a significance value of 0.05 was used. For every analysis, the strength of p values is reported in the figures according the following: p > 0.05, *p ≤ 0.05, **p ≤ 0.01, ***p ≤ 0.001, ****p ≤ 0.0001. Details of statistical tests can be found in the figure legends

## Acknowledgements

This work was supported by the Intestinal Stem Cell Consortium (U01DK103141 to JRS), a collaborative research project funded by the National Institute of Diabetes and Digestive and Kidney Diseases (NIDDK) and the National Institute of Allergy and Infectious Diseases (NIAID). This work was also supported by the NIAID Novel Alternative Model Systems for Enteric Diseases (NAMSED) consortium (U19AI116482 to JRS and ST). The content is solely the responsibility of the authors and does not necessarily represent the official views of the National Institutes of Health.

## Author contributions

Project conceptualization: MMC, SH, ST, JRS

Experimental design: MMC, JRS, MC, SH, YS, EA, MH, AJP

Experiments and data collection: MMC, MC, SH, YS, YHT, AW, BJ, NS, MH

Data analysis and interpretation: MMC, MC, SH, YHT, AW, MSN, BJ, NS, ST, MH, AJP, JRS

Writing manuscript: MMC and JRS

Editing manuscript: all authors

## References

Arora, N., Alsous, J.I., Guggenheim, J.W., Mak, M., Munera, J., Wells, J.M., Kamm, R.D., Asada, H.H., Shvartsman, S.Y., Griffith, L.G., 2017. A process engineering approach to increase organoid yield. Development 144, 1128–1136.

Augst, A.D., Kong, H.J., Mooney, D.J., 2006. Alginate Hydrogels as Biomaterials. Macromolecular Bioscience 6, 623–633.

Aurora, M., Spence, J.R., 2016. hPSC-derived lung and intestinal organoids as models of human fetal tissue. Dev Biol. 420, 230–238.

Bell, S., Schreiner, C., Wert, S., Mucenski, M., Scott, W., Whitsett, J., 2008. R-spondin 2 is required for normal laryngeal-tracheal, lung and limb morphogenesis. Development 135, 1049–1058.

Bray, N.L., Pimental, H., Melsted, P., Pachter, L., 2016. Near-optimal probabilistic RNA-seq quantification. Nature Biotechnology 34, 525–527.

Chaudhuri, O., 2017. Viscoelastic hydrogels for 3D cell culture. Biomaterials Science 5, 1480–2490.

Cruz-Acuna, R., Quiros, M., Farkas, A.E., Dedhia, P.H., Huang, S., Siuda, D., Garcia-Hernandez, V., Miller, A.J., Spence, J.R., Nusrat, A., Garcia, A.J., 2017. Synthetic hydrogels for human intestinal organoid generation and colonic wound repair. Nature Cell Biology 19, 1326–1335.

Czerwinski, M., Spence, J.R., 2017. Hacking the Matrix. Cell Stem Cell 20, 9–10.

Dame, M., Attili, D., McClintock, S., Dedhia, P.H., Ouillette, P., Hardt, O., Chin, A.M., Xue, X., Laliberte, J., Katz, E., Newsome, G., Hill, D.R., Miller, A.J., Tsai, Y.-H., Agorku, D., Altheim, C.H., Boscio, A., Simon, B., Samuelson, L., Stoerker, J., Appelman, H., Varani, J., Wicha, M., Brenner, D., Shah, Y., Spence, J.R., Colacino, J., 2018. Identificaion, isolation and characterization of human LGR5-positive colon adenoma cells. Development 145.

Dedhia, P.H., Bertaux-Skeirik, N., Zavros, Y., Spence, J.R., 2016. Organoid Models of Human Gastrointestinal Development and Disease. Gastroenterology 150, 1098–1112.

Dye, B., Dedhia, P.H., Miller, A.J., Nagy, M.S., White, E.S., Shea, L., Spence, J.R., 2016. A bioengineered niche promotes in vivo engraftment and maturation of pluripotent stem cell derived human lung organoids. eLife.

Dye, B., Hill, D.R., Ferguson, M., Tsai, Y.-H., Nagy, M.S., Dyal, R., Wells, J.M., Mayhew, C.N., Nattiv, R., Klein, O.D., White, E.S., Deutsch, G., Spence, J.R., 2015. In vitro generation of human pluripotent stem cell derived lung organoids. eLife.

Eiraku, M., Takata, N., Kawada, M., Sakakura, E., Okuda, S., Sekiguchi, K., Adachi, T., Sasai, Y., 2011. Self-organizing optic-cup morphogenesis in three-dimensional culture. Nature 472, 51–56.

Enemchukwu, N.O., Cruz-Acuna, R., Bongiorno, T., Johnson, C.T., Garcia, J.R., Sulchek, T., Garcia, A.J., 2016. Synthetic matrices reveal contributions of ECM biophysical and biochemical properties to epithelial morphogenesis. The Journal of Cell Biology 212, 113–124.

Finkbeiner, S.R., Freeman, J.J., Wieck, M.M., El-Nachef, W., Altheim, C.H., Tsai, Y.-H., Huang, S., Dyal, R., White, E.S., Grikscheit, T.C., Teitelbaum, D.H., Spence, J.R., 2015a. Generation of tissue-engineered small intestine using embryonic stem cell-derived human intestinal organoids. Biology Open 4, 1462–1472.

Finkbeiner, S.R., Hill, D.R., Altheim, C.H., Dedhia, P.H., Taylor, M.J., Tsai, Y.-H., Chin, A.M., Mahe, M.M., Watson, C.L., Freeman, J.J., Nattiv, R., Thomson, M., Klein, O.D., Shroyer, N.F., Helmrath, M.A., Teitelbaum, D.H., Dempsey, P.J., Spence, J.R., 2015b. Transcriptome-wide Analysis Reveals Hallmarks of Human Intestine Development and Maturation In Vitro and In Vivo. Stem Cell Reports 4, 1140–1155.

Forgacs, G., Foty, R.A., Shafrir, Y., Steinberg, M.S., 1998. Viscoelastic Properties of Living Embryonic Tissue: a Quantitative Study. Biophysical Journal 74, 2227–2234.

Fu, S., Thacker, A., Sperger, D.M., Boni, R.L., Valankar, S., Munson, E.J., Block, L.H., 2010. Rheological Evaluation of Inter-grade and Inter-batch Variability of Sodium Alginate. AAPS PharmSciTech 11, 1662–1674.

Gjorevski, N., Sachs, N., Manfrin, A., Giger, S., Bragina, M.E., Ordonez-Moran, P., Clevers, H., Lutolf, M.P., 2016. Designer matrices for intestinal stem cell and organoid culture. Nature 539, 560–564.

Heijmans, J., van Lidth de Jeude, J., Koo, B., Rosekrans, S., Wielenga, M., van de Wetering, M., Ferrante, M., Lee, A., Onderwater, J., Paton, J., Paton, A., Mommaas, A., Kodach, L., Hardwick, J., Hommes, D., Clevers, H., Muncan, V., van den Brink, G., 2013. ER stress causes rapid loss of intestinal epithelial stemness through activation of the unfolded protein response. Cell Rep 3, 1128–1139.

Hill, D.R., Huang, S., Nagy, M.S., Yadagiri, V.K., Fields, C., Muckherjee, D., Bons, B., Dedhia, P.H., Chin, A.M., Tsai, Y.-H., Thodla, S., Schmidt, T.M., Walk, S., Young, V.B., Spence, J.R., 2017. Bacterial colonization stimulates a complex physiological response in the immature human intestinal epithelium. eLife 6.

Huch, M., Knoblich, J.A., Lutolf, M.P., Martinez-Arias, A., 2017. The hope and hype of organoid research. Development 144, 938–941.

Jeon, O., Alt, D.S., Linderman, S.W., Alsberg, E., 2013. Biochemical and Physical Signal Gradients in Hydrogels to Control Stem Cell Behavior. Advanced Materials 25, 6366–6372.

Johnston, D.S., 2015. The Renaissance of Developmental Biology. PLoS Biology.

Josephson, R., Sykes, G., Liu, Y., Ording, C., Xu, W., Zeng, X., Shin, S., Loring, J., Maitra, A., Rao, M.S., Auerbach, J.M., 2006. A molecular scheme for improved characterization of human embryonic stem cell lines. BMC Bioloy 4.

Lancaster, M., Renner, M., Martin, C., Wenzel, D., Bicknell, L., Hurles, M., Homfay, T., Penninger, J., Jackson, A., Knoblich, J., 2013. Cerebral organoids model human brain development and microcephaly. Nature 501, 373–379.

Lee, K.Y., Mooney, D.J., 2012. Alginate: properties and biomedical applications. Prog Polym Sci. 37, 106–126.

Leslie, J., Huang, S., Opp, J., Nagy, M.S., Kobayashi, M., Young, V.B., Spence, J.R., 2015. Persistence and toxin production by Clostridium difficile within human intestinal organoids result in disruption of epithelial paracellular barrier function. Infection and Immunity 83, 138–145.

Little, M.H., 2017. Organoids: a Special Issue. Development 144, 935–937.

Love, M.I., Huber, W., Anders, S., 2014. Moderated estimation of fold change and dispersion for RNA-seq with DESeq2. Genome Biology 15:550.

McCracken, K., Howell, J., Wells, J.M., Spence, J.R., 2011. Generating human intestinal tissue from pluripotent stem cells in vitro. Nature Protocols 6, 1920–1928.

Miller, A.J., Hill, D.R., Nagy, M.S., Aoki, Y., Dye, B., Chin, A.M., Huang, S., Zhu, F., White, E.S., Lama, V., Spence, J.R., 2018. In Vitro Induction and In Vivo Engraftment of Lung Bud Tip Progenitor Cells Derived from Human Pluripotent Stem Cells. Stem Cell Reports 10, 101–119.

Munera, J., Sundaram, N., Rankin, S., Hill, D., Watson, C., Mahe, M., Vallance, J., Shroyer, N., Sinagoga, K., Zarzoso-Lacoste, A., Hudson, J., Howell, J., Chatuvedi, P., Spence, J., Shannon, J., Zorn, A., Helmrath, M., Wells, J., 2017. Differentiation of Human Pluripotent STem Cells into Colonic Organoids via Transient Activation of BMP Signaling. Cell Stem Cell 21, 51–64.

Rowley, J.A., Madlambayan, G., Mooney, D.J., 1999. Alginate hydrogels as synthetic extracellular matrix materials. Biomaterials 20, 45–53.

Samorezov, J.E., Morlock, C.M., Alsberg, E., 2015. Dual Ionic and Photo-Crosslinked Alginate Hydrogels for Micropatterned Spatial Control of Material Properties and Cell Behavior. Bioconjugate Chemistry 26, 1339–1347.

Singagoga, K.L., Wells, J.M., 2015. Generating human intestinal tissues from pluripotent stem cells to study development and disease. The EMBO Journal 34, 1149–1163.

Spence, J.R., Mayhew, C.N., Rankin, S.A., Kuhar, M., Vallance, J.E., Tolle, K., Hoskins, E.E., Kalinichenko, V.V., Wells, S.I., Zorn, A.M., Shroyer, N.F., Wells, J.M., 2011. Directed differentiation of human pluripotent stem cells into intestinal tissue in vitro. Nature 470, 105–109.

Takasato, M., Er, P.X., Becroft, M., Vanslambrouck, J.M., Stanley, E.G., Elefanty, A.G., Little, M.H., 2014. Directing human embryonic stem cell differentiation towards a renal-lineage generates a self-organizing kidney. Nature Cell Biology 16, 118–126.

Takebe, T., Sekine, K., Enomura, M., Koike, H., Kimura, M., Ogaeri, T., Zhang, R.-R., Ueno, Y., Zheng, Y.-W., Koike, N., Aoyama, S., Adachi, Y., Taniguchi, H., 2013. Vascularized and functional human liver from an iPSC-derived organ bud transplant. Nature 499, 481–484.

Tsai, Y.-H., Czerwinski, M., Wu, A., Dame, M., Attili, D., Hill, E., Colacino, J., Nowacki, L.M., Shroyer, N., Higgins, P.D., Kao, J.Y., Spence, J.R., 2018. A Method for Cryogenic Preservation of Human Biopsies and Subsequent Organoid Culture. Cellular and Molecular Gastroenterology and Hepatology.

Tsai, Y.-H., Hill, D.R., Kumar, N., Huang, S., Chin, A.M., Dye, B., Nagy, M.S., Verzi, M., Spence, J.R., 2016. LGR4 and LGR5 Function Redundantly During Human Endoderm Differentiation. Cel Mol Gastroenterol Hepatol. 2, 648–662.

Tsai, Y.-H., Nattiv, R., Dedhia, P.H., Nagy, M.S., Chin, A.M., Thomson, M., Klein, O.D., Spence, J.R., 2017. In vitro patterning of pluripotent stem cell-derived intestine recapitulates in vivo human development. Development 144, 1045–1055.

Watson, C., Mahe, M., Junera, J., Howell, J., Sundaram, N., Poling, H., Schweitzer, J., Vallance, J., Mayhew, C., Sun, Y., Grabowski, G., Finkbeiner, S.R., Spence, J.R., Shroyer, N., Wells, J.M., Helmrath, M., 2014. An in vivo model of human small intestine using pluripotent stem cells. 2014 20, 1310–1314.

Webber, R.E., Shull, K.R., 2004. Strain dependence of the viscoelastic properties of alginate hydrogels. Macromolecules 37, 6153–6160.

Wells, J.M., Spence, J.R., 2014. How to make an intestine. Development 141, 752–760.

